# Exaggerated in vivo IL-17 responses discriminate recall responses in active TB

**DOI:** 10.1101/516690

**Authors:** Gabriele Pollara, Carolin T Turner, Gillian S Tomlinson, Lucy CK Bell, Ayesha Khan, Luis Felipe Peralta, Anna Folino, Ayse Akarca, Cristina Venturini, Tina Baker, Fabio LM Ricciardolo, Teresa Marafioti, Cesar Ugarte-Gil, David AJ Moore, Benjamin M Chain, Mahdad Noursadeghi

## Abstract

Host immune responses at the site of *Mycobacterium tuberculosis* (Mtb) infection serve to contain the pathogen, but also mediate the pathogenesis of tuberculosis (TB) and onward transmission of infection. Interferon gamma (IFNγ) responses do not discriminate between protection and pathogenicity, but IL-17A/F responses, known to drive pathology in diverse chronic inflammatory diseases, have also been associated with TB pathogenesis in animal models. At the site of in vivo immune recall responses to Mtb modelled by the tuberculin skin test, we show for the first time that active TB in humans is also associated with exaggerated IL-17A/F expression, accumulation of Th17 cells and IL-17A/F bioactivity, including increased neutrophil recruitment and matrix metalloproteinase-1 expression directly implicated in TB pathogenesis. These features discriminate recall responses in patients with active TB from those with cured or latent infection and are also evident at the site of TB disease. Our data support targeting of this pathway in host-directed therapy for TB.

## Introduction

*Mycobacterium tuberculosis* (Mtb) infection results in a spectrum of clinical outcomes, from asymptomatic latent infection to symptomatic disease. The focus of host-pathogen interactions is characterised histologically by granuloma formation, a chronic inflammatory process that may contain the infection, but can also result in tissue damage that promotes transmission of infection to other individuals (1, 2). The distinctions that tip the balance between protective and pathogenic immune responses remain a fundamental question in tuberculosis research. This knowledge is expected to inform rational vaccine design and development of host-directed therapies (3).

**Figure 1.**
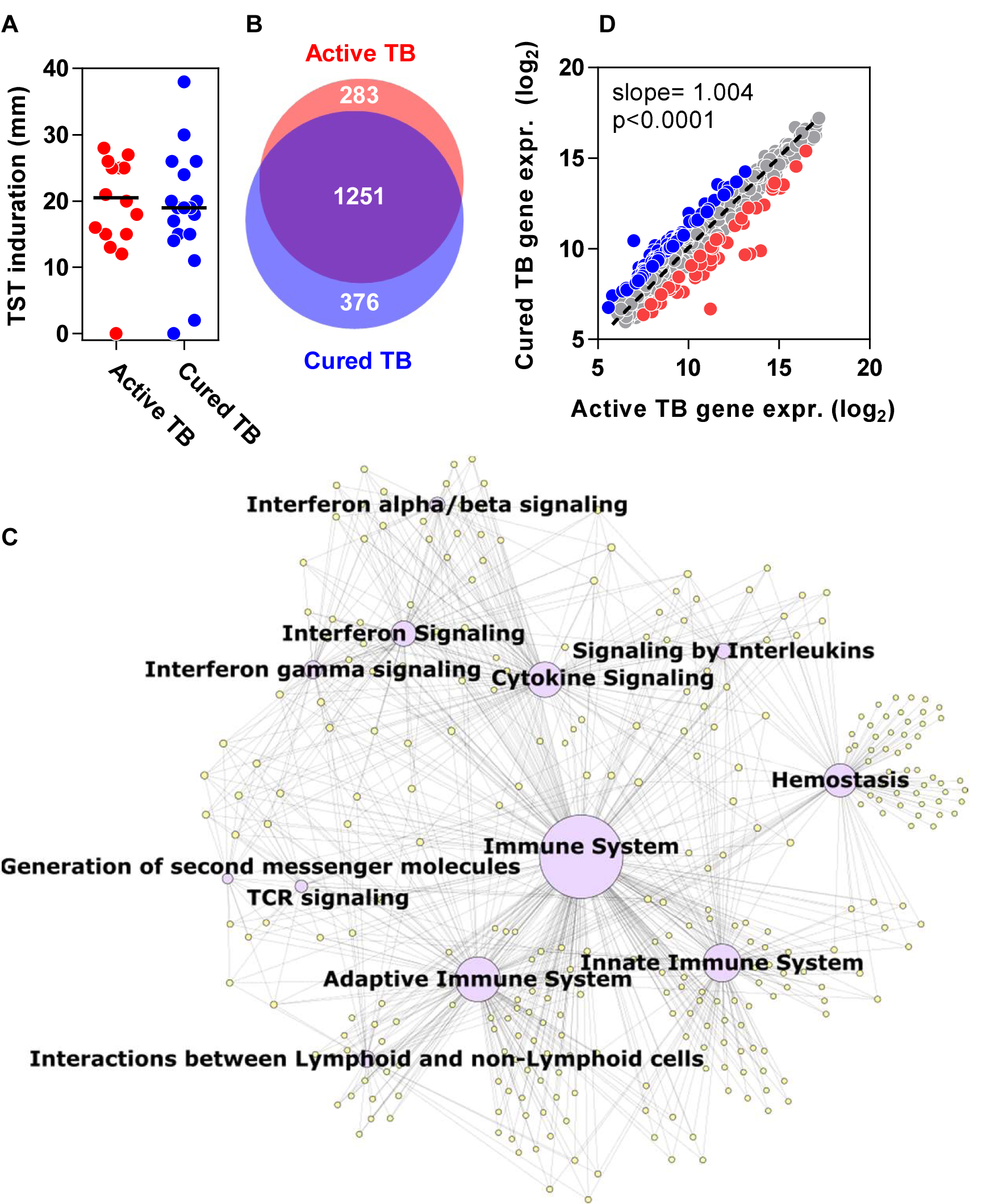
TST transcriptome in patients with active and cured TB disease. A) Induration at the site of TST was recorded by routine clinical assessments in both populations (mm) and was not different between the two groups (Mann-Whitney test). B) Venn diagram depicting genes significantly upregulated 2-fold in the TST of patients with active or cured TB relative to control saline injection. (p<0.01 by t-test with no multiple testing correction). C) TST transcriptome common to both active and cured TB summarised as a network diagram. Purple nodes represent Reactome database functional pathways, yellow nodes represent genes and edges reflect relationship between pathways and genes. Pathway node diameters are proportional to the respective pathway –log10 p value enrichment statistic. D) Pairwise dot-plot of 1910 gene integrated TST signature in patients with either active or cured TB. Dotted line reflects line of perfect covariance. p value derived from linear regression model between the two variables. Red and blue dots represent genes >2 fold differentially expressed (p<0.01 by t-test with no multiple testing correction) between active and cured TB.

Chronic inflammatory pathology at the site of human tuberculosis has been the subject of extensive descriptive studies, but discriminating between protective and pathological immune responses has been limited to comparing leucocyte phenotype and function in blood from Mtb-exposed patients with and without active disease (1). We have shown that genome-wide transcriptional profiling of biopsies of the tuberculin skin test (TST) can be used to make comprehensive molecular and systems level assessment of in vivo immune responses at the site of standardised host-pathogen interactions (4–7). Importantly, the transcripts enriched within the TST reflect the genome-wide variation in molecular pathology at the site of tuberculosis (TB) disease (5, 7), suggesting the TST represents a valuable surrogate for assessing TB immunopathogenesis in vivo.

On the premise that active TB disease is predominantly a manifestation of immunopathology, in this study we aimed to test the hypothesis that immune responses at the site of host-pathogen interactions, modelled by the TST, would reveal enrichment of immunopathologic responses in patients with active TB that were absent in individuals with equivalent immune memory for Mtb but without disease.

## Results

### Immune responses at the site of TST in active and cured TB

The TST has been most extensively used to identify patients with T cell memory for mycobacterial antigens, but the clinical response does not differentiate infected individuals with and without active disease (1). We sought to test the hypothesis that molecular profiling of the TST may identify elements of the recall response which are specifically associated with disease. We undertook 48-hour TSTs in patients with microbiologically-confirmed TB disease within the first month of treatment (‘active TB’) to identify disease associated responses, and compared these to TST responses in patients within one year of curative TB treatment (‘cured TB’) to identify non-disease associated recall responses (table E1). Age, gender and site of TB disease were comparable between the two groups (table E2). As expected, clinical induration in response to the TST was not significantly different between the two groups (fig 1A), hitherto interpreted to reflect comparable cell mediated immune memory.

In comparison to skin biopsies from the site of control saline injection, 1910 genes were significantly enriched in response to the TST in at least one study group. Of these, 1251 were enriched in both groups (fig 1B). Bioinformatic systems level assessment of the shared response revealed many prototypic cell mediated immune responses which we had previously described in the TST (fig 1C) (5). Pairwise assessment of the integrated list of transcripts that were enriched in either group revealed statistically significant covariance, consistent with the hypothesis that the majority of responses do not discriminate between the two groups (fig 1D).

### Differential gene expression in the TST in active TB

A proportion of genes were differentially enriched between the two groups (fig 1D & tables E3-4). 44 genes were expressed significantly more in patients with active TB (table E3) compared to patients with cured TB. Amongst these, pathway analysis identified statistically significant enrichment of transcripts involved in extracellular matrix (ECM) remodelling, such as matrix metalloproteinase 1 (MMP-1), and beta defensins that both exert antimicrobial functions and also provide a chemotactic gradient for CCR2-expressing cells, including neutrophils (8) (figs 2A & E1A). MMP-1, previously implicated in pathogenic degradation of the ECM in TB (9), was the most over-expressed gene in active TB compared to cured TB (fig 2B & table E3). This difference was validated at protein level by immunohistochemistry, which also revealed that the differences in MMP-1 expression between patients with active and cured TB was restricted to the inflammatory infiltrates within the TST (fig 2C-E).

**Figure 2.**
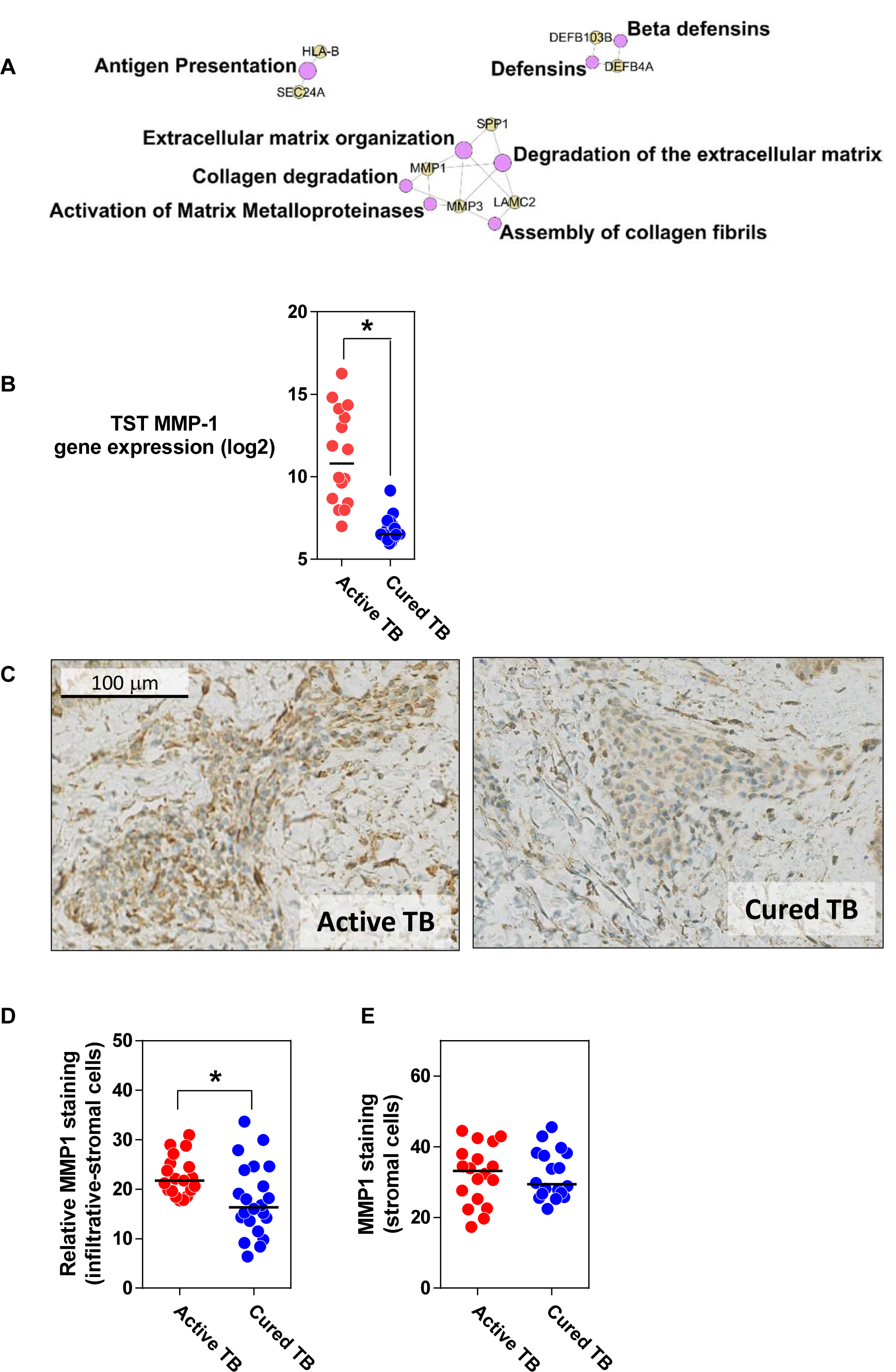
MMP-1 over-expression in active TB. A) Network diagram of genes and Reactome pathways over-expressed in the TST of patients with active TB compared to cured TB. Purple nodes represent Reactome database functional pathways, yellow nodes represent genes and edges reflect relationship between pathways and genes. Pathway node diameters are proportional to the respective pathway – log10 p value enrichment statistic. B) mRNA expression in TST of patients with active and cured TB of MMP1 gene. C) Representative MMP-1 immunohistochemistry staining in inflammatory infiltrates with TST samples from patients with active and cured TB. D) Differential MMP-1 staining intensity for 18 cellular infiltrates in each group of patients relative to adjacent zones of skin with no cellular infiltration. E) MMP-1 staining intensity in TST zones outside inflammatory infiltrates. * = p<0.01 by Mann-Whitney test.

### Elevated IL-17 responses in active TB

We hypothesised that the genes over-expressed in active TB were regulated by common upstream signals in the tissue environment. To test this hypothesis, we compared the predicted upstream regulators of differentially expressed transcripts in the TST of active and cured TB patients using Ingenuity Pathway Analysis (6). This analysis suggested that IL-17A induced the expression of genes over-expressed in active TB and not those over-expressed in cured TB (fig 3A). In contrast, IFNγ was predicted to be an upstream signal for gene expression enriched in both active and cured TB (fig 3A). IFNγ responses, largely attributed to T helper (Th)-1 polarised CD4+ T cells are necessary for immunological protection against Mtb (10), but they are insufficient and do not discriminate between people who do and do not develop disease (1). The role of IL-17A in TB is less clear. This cytokine belongs to a family of six structurally related cytokines and shares greatest sequence homology with IL-17F. These bind the same receptor, and consequently exert the same functions, particularly increased neutrophil recruitment via upregulation of chemokine expression (11).

**Figure 3.**
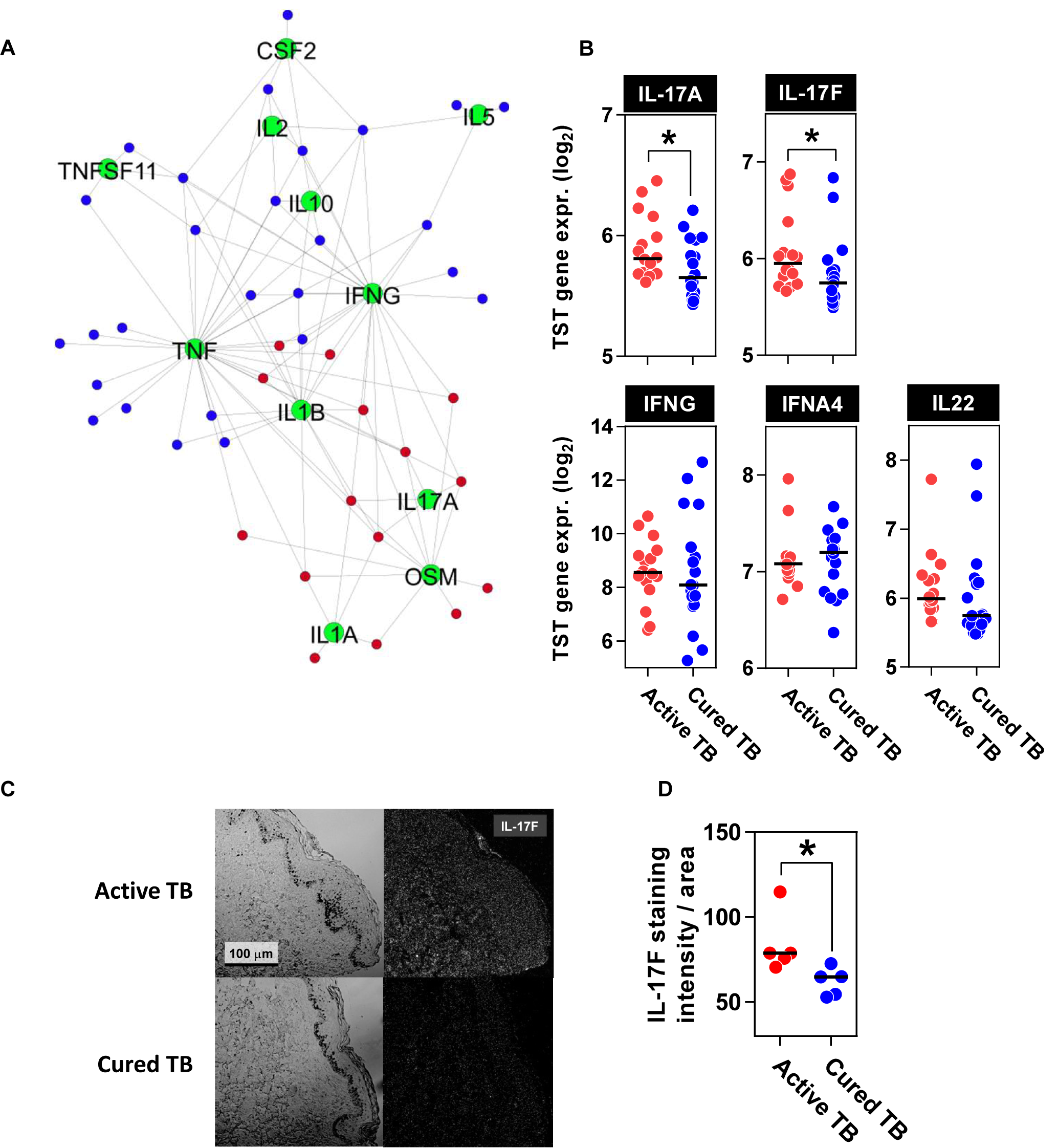
IL-17 is over expressed in active TB. A) Network diagram depicting upstream cytokine analysis of genes differentially expressed in TST of patients with active and cured TB. Red and blue nodes represent significant genes overexpressed in active and cured TB respectively as identified in fig 1D. Green nodes represent predicted cytokines regulating the gene expression of red and blue nodes. Edges depict relationship between upstream regulators and differentially expressed genes. B) Expression of selected cytokine transcript within the TSTs of active and cured TB patients. C) Expression of IL-17F by immunofluorescence in TST of patients with active TB. Left panel = phase contrast image, right panel = IL-17F positivity (white). D) Quantification of IL-17F staining throughout TST sections from patients with active and cured TB, determined by pixel intensity as a proportion of area sampled. Each dot represents IL-17F expression in the cross-section of an entire TST biopsy from one patient. * p<0.01 by Mann-Whitney test.

There is unequivocal evidence that IL-17A/F contribute to host defence against bacterial and fungal pathogens (12). Importantly however, they are also strongly indicated in the immunopathology of chronic inflammatory diseases (12, 13). This includes evidence for IL-17A dependent neutrophil mediated pathology in mouse models of Mtb infection (14–16). Our bioinformatics analysis suggested increased enrichment of transcripts in the human in vivo recall response of patients with active TB may be driven by IL-17A/F activity. Therefore, we sought to test the hypothesis that IL-17A/F activity is exaggerated in active TB. Consistent with this hypothesis the expression of both IL-17A and IL-17F were enriched in the TST of people with active TB compared to cured TB (fig 3B). In contrast, IFNγ transcript levels representing the prototypic molecule in cell mediated immune recall responses was not significantly different (fig 3B). Interestingly, the expression of IL-22, a cytokine with closely related biological function to IL-17A and IL-17F (17), was also not elevated in active TB (fig 3B). The differences revealed at the transcriptional level were also reflected by increased immunofluorescence of IL-17F protein in the TST of people with active TB (fig 3C & 3D).

In order to test the functional significance of the differences in IL-17A/F expression between active and cured TB, we evaluated differences in the bioactivity of IL-17A between these groups. We addressed this by generating cellular response modules from the transcriptomes of cytokine-stimulated keratinocytes (KC) (18, 19). We confirmed that these modules were both sensitive and specific for their cognate stimuli by assessing their expression in other independent datasets (fig E2 & table E6). We then compared the geometric mean expression of these cytokine-specific transcriptional modules in the TST transcriptomes. The IL-17A-induced gene module was significantly increased in the TST of people with active TB compared to that of cured TB, but expression of IFNγ, type I IFN or TNF-inducible gene modules was not significantly different between the two groups (fig 4A). We extended our approach to evaluating the functional bioactivity of other specific cytokines using transcriptional modules for IL-10-and IL-4/IL-13-inducible gene expression, described in a previous report (5). Neither of these were significantly different in the TST of people with active and cured TB (fig E3A). The IL-4/IL-13 bioactivity module was used as a measure of Th2 responses normally associated with allergy and immune responses to helminths. Another member of the IL-17 family, IL-17E contributes to Th2 responses (11). We found no enrichment of IL-17E expression in the TST of people with active TB consistent with IL-4/IL-13 bioactivity and distinct from IL-17A/F bioactivity (fig E3B).

**Figure 4.**
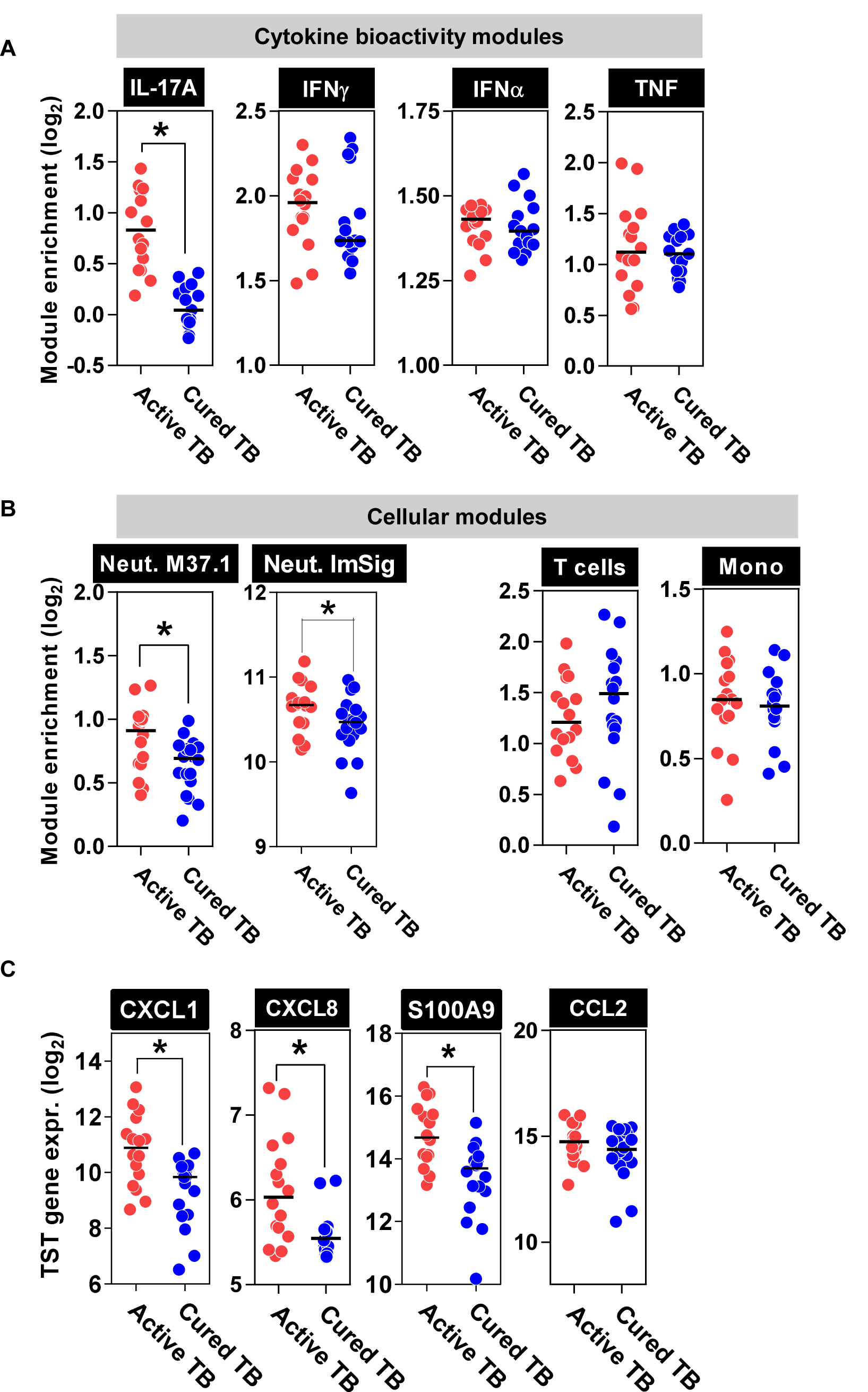
TST challenge in active TB is characterised by enrichment of IL-17 responses. Enrichment in TST relative to saline injection of A) keratinocyte response modules of cytokine bioactivity, B) immune cell modules and C) CXCL1, CXCL8, S100A9 and CCL2 genes. Neutrophil (Neut.) modules M37.1 and ImSig validated for sensitivity and specificity in references (23, 24) respectively. * = p<0.01 by Mann-Whitney test.

Focusing on mechanisms that may contribute to pathogenesis, enrichment of MMP-1 expression in the TST of patients with active TB (fig 2) can also be attributed to increased IL-17A/F bioactivity, through induction of MMP expression by stromal cells (20). Neutrophils can also contribute to the immunopathology of TB (14–16, 21, 22), and a key function of IL-17A/F is to promote neutrophil recruitment via induction of neutrophil chemokines (12). Therefore, we tested the hypothesis that the TST in active TB will also reveal increased neutrophil recruitment, compared to that of patients with cured TB. We compared the expression of two independently derived transcriptional modules that have been extensively validated to reflect neutrophil frequency in tissues including the TST (23, 24). We found significantly higher expression of the neutrophil modules in people with active TB compared to cured TB (fig 4B). These differences were mirrored by gene expression levels of IL-17A-inducible CXCL1, CXCL8 and S100A9 that drive neutrophil recruitment to sites of inflammation (11, 15) (fig 4C). In contrast, accumulation of monocytes and T cells assessed by their respective gene expression modules (23), and the expression of the monocyte chemoattractant, CCL2, did not differ in the TST of people with active and cured TB (fig 4B&C).

### Increased frequency of Th17 cells in active TB TST responses

IL-17A/F are predominantly produced by Th17 cells and neutrophils (11). Immunohistochemistry revealed that in the TST of people with active TB, IL-17F, originated predominantly from mononuclear cells, rather than from polymorphonuclear cells (fig 5A). Therefore, we tested the hypothesis that Th17 cells were enriched in the TST of people with active TB compared to that of cured TB. We derived transcriptional modules specific for differentially polarised CD4+ Th subsets from a published dataset (25) and demonstrated their specificity in an independent dataset (26) (fig E4A & table E7). We further validated the Th17 module by showing that this correlated closely with the expression of the IL-17A/F bioactivity module within skin biopsies of patients with psoriasis vulgaris, representing an alternative Th17-mediated inflammatory condition (27) (fig E4B). The expression of the Th1- and Th2-associated transcriptional modules in the TST was comparable between patients with active and cured TB, but expression of the Th17 associated module was significantly increased in patients with active TB (figs 5B & E3).

**Figure 5.**
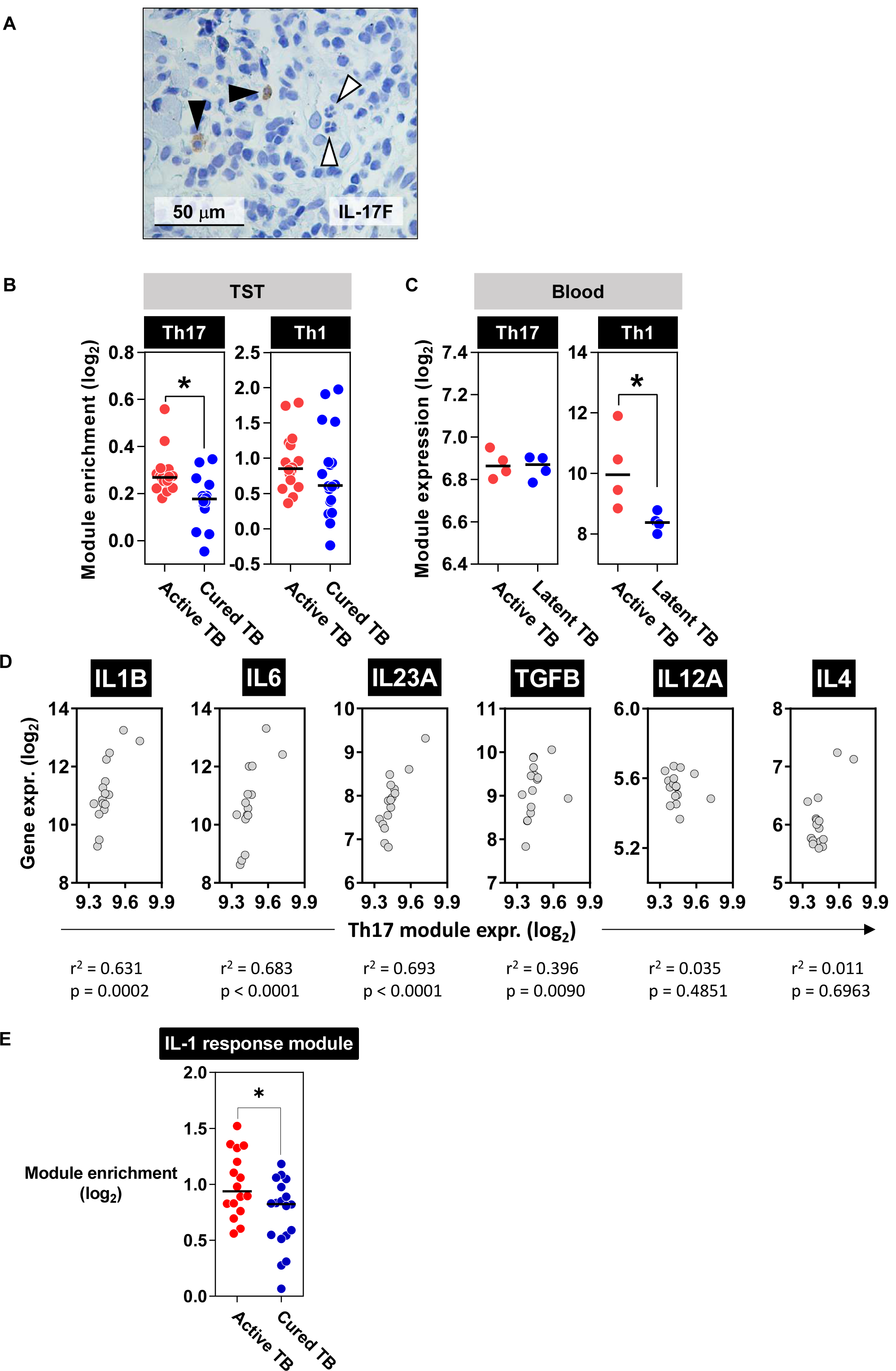
Active TB is characterised by elevated Th17 cells. A) IL-17F immunohistochemistry in TST of patients with active TB. Black arrows point to mononuclear cells that express IL-17F and white arrows point to polymorphonuclear cells that do not express IL-17F. B) Th17 and Th1 cell module enrichment in TST relative to saline injection. C) Expression of Th17 and Th1 modules in PPD-stimulated PBMC from patients with active or latent TB (data originated from dataset GSE27984). * = p<0.01 by Mann-Whitney test. D) Relationship between expression of Th17 transcriptional module and individual cytokine gene expression in TST of cytokines implicated in polarising CD4+ T cells to a Th17 phenotype. Data points represent patients with active TB. Spearman rank correlation coefficients (r^2^) and p values. D) Enrichment in TST relative to saline injection of IL-1β fibroblast response modules indicative of IL-1 cytokine bioactivity. * = p<0.01 by Mann-Whitney test.

In order to investigate the mechanism for increased Th17 responses in the TST of patients with active TB, we tested the hypothesis that active TB was also associated with increased circulating Th17 cells. We assessed the expression of the Th subset modules in the transcriptome of PPD-stimulated PBMC from an independent cohort of individuals with active TB disease and latent TB infection (28). In contrast to that observed in the TST, patients with active TB revealed enrichment for the Th1-associated gene expression module in PBMC but showed no difference in the Th17-associated module (fig 5C). As a result, we explored the alternative hypothesis that the inflammatory environment generated at the site of TST promotes the Th17 differentiation observed in patients with active TB. We investigated the expression of cytokines implicated in Th17 differentiation, IL-1β, IL-6, IL-23 and TGFβ (11). The expression of each of these was significantly correlated with enrichment for Th17 cells in the TST of active TB. In contrast there was no correlation with IL-12A and IL-4 expression that drive Th1 and Th2 cell differentiation respectively (fig 5D). To test the hypothesis that conditions favouring Th17 differentiation were present in vivo, we generated and validated a transcriptional response module for IL-1 activity in human fibroblasts (fig E5 and table E8) in order to quantify functional IL-1 activity in the TST. This analysis confirmed increased expression of IL-1 inducible genes in the TST of patients with active TB (fig 5E). Of note, this finding was consistent with the bioinformatic prediction of IL1 signalling upstream of some transcripts that were differentially enriched predominantly in active TB (fig 3A). Overall, these data support a model in which the local tissue microenvironment, in part driven by elevated IL-1 activity, may promote Th17 differentiation within tissues in active TB.

### IL-17 activity is not a feature of latent TB and is not confounded by demographic background, extrapulmonary disease or time on treatment

In order to validate our findings, we compared the TST transcriptome of a second independent cohort of people with active TB with that of individuals with latent TB infection. Consistent with our previous results, this comparison revealed significant enrichment for IL-17A/F bioactivity and neutrophil infiltration in people with active TB (fig 6A). Furthermore, Th17 cells were enriched in those with active TB despite there being no overall difference in T cell numbers compared to the TST in individuals with latent TB (fig 6A). Of note, the comparison to latent TB in this analysis, also excluded the possibility that lower levels of IL-17A/F bioactivity in the TST of people with cured TB compared to active TB was an off-target effect of the antimicrobial treatment for active TB.

**Figure 6.**
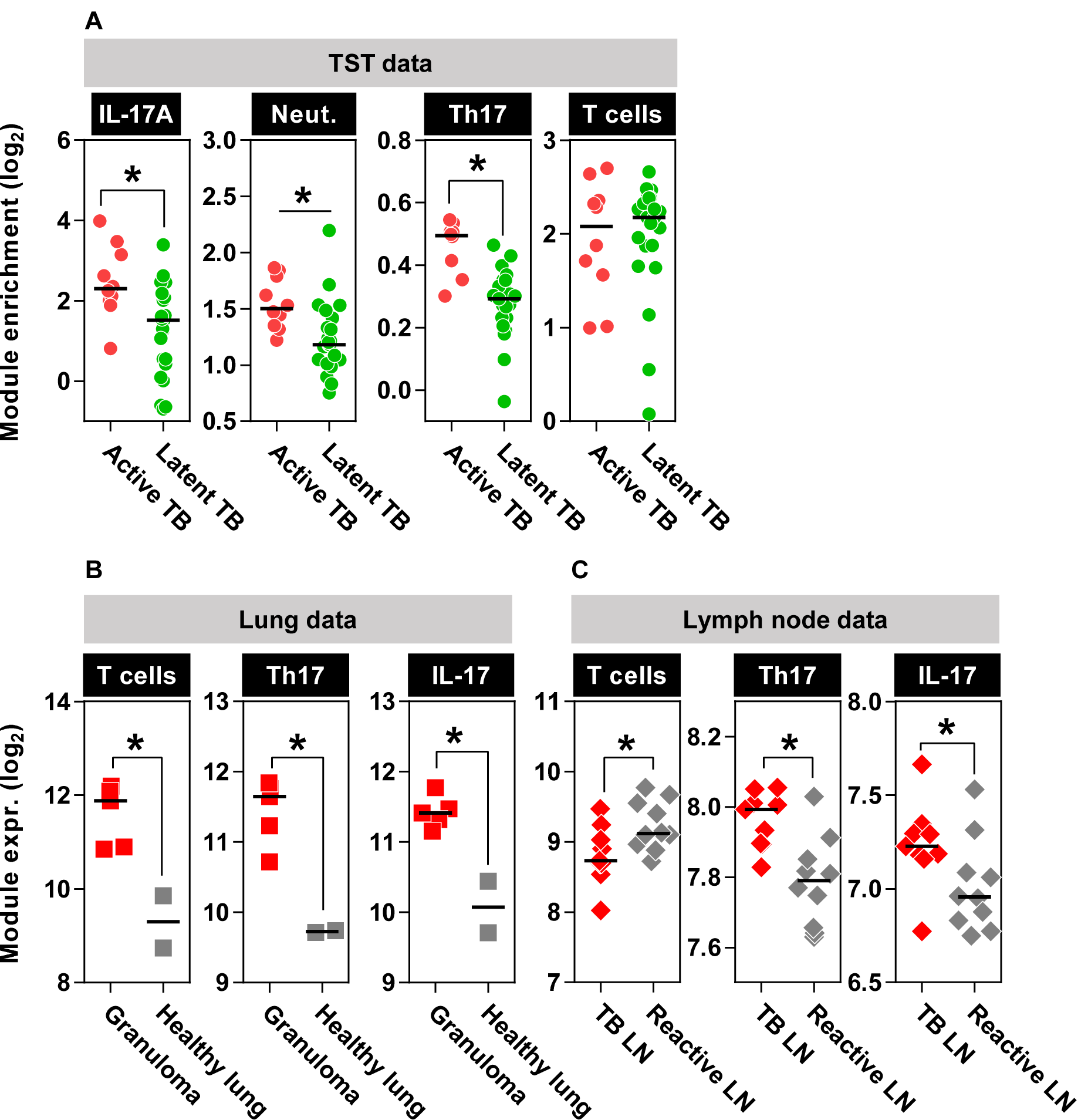
Th17 cells and IL-17 activity are a feature of both TST reactions and sites of human TB disease. A) Enrichment in TST relative to saline injection of transcriptional modules for IL-17 bioactivity, neutrophils, Th17 cells and T cells in TST of independent cohort of patients with active TB and a separate cohort of individuals with latent TB infection. B) Expression of T cells, Th17 cells and IL-17 bioactivity modules from the site of human TB granuloma relative to healthy lung tissue (dataset GSE20050) and C) in human TB infected lymph nodes (LN) relative to reactive lymph nodes that do not display evidence of granulomatous inflammation or cancer (dataset E-MTAB-2547). * = p<0.01 by Mann-Whitney test.

Taken together our data show that active TB is associated with exaggerated Th17 recall responses and IL-17 bioactivity within the tissue microenvironment of host-pathogen interactions. Our findings were replicated using two different methods for transcriptional profiling (microarray figs 4-5 and RNA-Seq fig 6), and were consistent in geographically distinct cohorts (fig E6A). Significantly exaggerated Th17 recall and IL-17A/F bioactivity in people with active TB compared to either cured or latent TB were preserved in analyses including only UK cohorts (figs E6B & E6C), confirming that the differences were not due to confounding by other variables in the different cohorts. In addition, we found no difference in these responses between people with pulmonary and extrapulmonary TB disease (fig E7). Of note, people with active TB were assessed at different time points within the first month of treatment (median 11 days, IQR 5-28 days). Rapid changes in the peripheral blood transcriptome associated with active TB have been reported (29, 30), but we found no diminution of the exaggerated Th17 recall responses and IL-17 bioactivity invoked by the TST challenge in this time frame (fig E8), suggesting that the mechanisms that drive these responses in active are not swiftly reversed by treatment.

### IL-17 activity is present at the site of TB disease

Having established that IL-17A/F production by T cells was a prominent feature of the in vivo tissue recall response to Mtb stimulation, we sought to determine whether this was also evident at the site of TB disease. The transcriptome of human TB granuloma (31) showed an enrichment of T cells, Th17 cells and IL-17A bioactivity compared to normal lung tissue (fig 6B). In addition, we examined the transcriptome of human Mtb-infected lymphadenitis compared to other ‘reactive’ causes of lymphadenopathy devoid of granulomatous inflammation or malignancy (32). Interestingly, despite the fact that reactive lymph nodes were enriched for the T cell transcriptional module, the Th17 and IL-17A/F bioactivity modules were enriched within Mtb infected lymph nodes (fig 6C), further confirming enrichment for IL-17A/F bioactivity to be a feature at the site of TB disease.

## Discussion

Active TB disease is characterised by chronic inflammation that can result in significant tissue destruction, necessary for the onward transmission of Mtb (2). Identifying the processes that govern this immunopathology offers the opportunity to intervene therapeutically, limiting tissue damage and transmission of infection. Pathogenic immune pathways have been difficult to identify because most components of the immune response to Mtb do not discriminate between different clinical outcomes of infection (1). The present study provides compelling new evidence that IL-17A/F responses may mediate immunopathology in active TB. The inclusion of multiple cohorts and diverse demographic backgrounds increased the generalisability of our findings and circumvented the limitations of our cross-sectional study design. The findings are consistent with the well-established role of IL-17A/F in the immunopathology of chronic inflammatory human disease exemplified by psoriasis, for which blockade of the IL-17A/F axis provides an effective treatment (13).

Importantly, the primary physiological role of IL-17A/F is to promote host defence against bacterial and fungal infection, cogently demonstrated by mice deficient for IL-17A/F or IL-17 receptors, and by humans with inborn errors of IL-17 immunity (12). Mouse models also suggest a protective role for IL-17A/F following BCG vaccination (33) and in the early stages of Mtb infection, particularly in the context of more virulent Mtb strains (34, 35), through localising T cells near Mtb-infected macrophages (34) and by preventing formation of necrotic granuloma (36). However, in mice rendered susceptible to TB disease as a result of IFNγ deficiency or through receiving repeated BCG vaccinations, IL-17A/F responses drive neutrophil-mediated pathology (14–16, 21). We infer from these data that IL-17A/F responses may play a dichotomous role in TB by contributing to protection in early infection, but to the pathology of disease if early infection is not controlled and multibacillary bacterial replication and chronic immune activation ensues. Additional support for this model is evident in IL27R deficient mice, which exhibit exaggerated Th17 responses because they lack IL-27 inhibition of RORγT (37). These mice show enhanced clearance of Mtb, but also increased immunopathology dependent on IL-17A. Interestingly, a human candidate gene study identified host genetic variation associated with increased secretion of IL-17A to be correlated with both protection from incident TB but more severe TB disease (38). Taken together, we hypothesise that exaggerated IL-17A/F recall responses are the consequence of chronic multibacillary infection, but in turn mediate increased pathology in established disease. Of the other IL-17 family members that signal via alternative receptors, much less is known about the functional role of IL-17B, IL-17C and IL-17D (11). These cytokine responses were not specifically tested in the present study. IL-17E, known to promote Th2 responses (11), was not differentially enriched in active TB. Equally, we found no evidence of elevated IL-10 responses in active TB, indicating an uncoupling of IL-17A/F activity from regulatory responses, driving immunopathology rather than the control of Mtb replication (39). Another cytokine, IL-22, does have functional overlap with IL-17A/F (17), but did not discriminate between people with and without disease in our study.

Our data support the hypothesis that exaggerated IL-17A/F responses arise from Th17 cells, but they do not unequivocally exclude other cellular sources. Our immunohistochemical analysis did not show any clear evidence for IL-17F production by polymorphonuclear cells in the TST, but we were not able to test whether other lymphocyte populations, such as γδ T cells, made a significant contribution (35, 40, 41). Future studies will require single cell resolution to definitively confirm the source of exaggerated IL-17A/F responses in this model, as well as to determine whether active TB shows enrichment for ‘pathogenic’ Th17 cells that express both IFNg and IL-17A/F (42). Consistent with previous studies (40, 43), we found no evidence of increased circulating Th17 cells in active TB, although this was measured in a separate active TB cohort to the ones that underwent TST assessment. Nevertheless, this observation underscores the unique value of molecular level assessment of immune responses in the tissue microenvironment at the site of host-pathogen interactions, modelled by the TST. Importantly, it also suggests the model that differential Th17 responses within the TST are governed by immune signalling networks within the tissue microenvironment after T cell recruitment. In active TB patients, the tissue expression of cytokines that promote Th17 differentiation correlated with the transcriptional module for Th17 cells, and elevated IL-1 activity was present in the TST. The source of these cytokines is likely to be myeloid cells (11, 13, 44). Although we cannot prove a causal relationship at this stage, our data are consistent with a model in which infiltrating monocytes cells from patients with active TB secrete elevated levels of cytokines that promote the differentiation of recruited T cells to a Th17 phenotype. Of note, active TB patients have higher frequency of circulating CD14+ CD16+ non-classical monocytes (45), which can potentiate the differentiation of CD4 T cells to a Th17 phenotype in patients with chronic inflammatory conditions (46).

Our findings challenge the long-established view that curative treatment of TB does not lead to contemporaneous changes to immunological recall responses (47). Our data are consistent with the hypothesis that IL-17A-inducible neutrophil chemotaxis and expression of MMP-1 represents a key mediator of immunopathology in TB by promoting granuloma formation, bacterial replication and matrix degradation that can lead to cavitation and onward transmission (9, 48). Our results support future studies to evaluate the impact of modulating IL-17A/F activity to ameliorate the pathology associated with chronic TB disease. The availability of therapies that block the IL-17A/F cytokine axis, developed and licensed for chronic inflammatory diseases (13), offers invaluable opportunities to transition from proof of concept pre-clinical studies, for example in non-human primate models, to first in man experiments.

## Methods

### Study populations

The cohorts of patients recruited to the study are described in detail in Supplementary Methods. Inclusion and exclusion criteria are in table E1 and the demographic, clinical and laboratory data for each study group is summarised in table E2.

### Study approval

Recruitment of patients with cured and latent TB was approved by UK National Research Ethics Committee (reference number: 14/LO/0505) and Universidad Peruana Cayetano Heredia Institutional Ethics Committee (reference number: 62349). Recruitment of patients for the validation active TB cohort was approved UK National Research Ethics Committee (reference number: 16/LO/0776).

### Study schedule and sampling

On recruitment to the study, all participants received 0.1 mL intradermal injection of two units tuberculin (Serum Statens Institute) or saline in the volar aspect of one forearm, and this site was marked with indelible ink. Measurement of the TST site, biopsy collection and sample storage was performed as previously described (4) and is expanded in more detail in Supplementary Methods.

### Sample processing

The TST transcriptome from active TB patients in the discovery cohort was derived directly from the data repository E-MTAB-3254 (ArrayExpress - https://www.ebi.ac.uk/arrayexpress/). Skin samples from all other participants were processed for RNA extraction and transcriptional profiling by either microarrays or RNA-Seq as described in Supplementary Methods.

### Whole genome transcriptional profiling and analysis software

Raw microarray data was processed and normalised as previously described (49). Raw RNASeq counts were normalised within-sample into TPM (transcripts per million) to remove feature-length and library-size effects (50). Log2 transformed TPM were used for further analysis, including transcriptional module derivation and expression that is described in further detail in Supplementary Methods.

### Transcriptomic data repositories

All transcriptional datasets used in this study are described in table E5. Accession numbers refer to datasets in the ArrayExpress repository (https://www.ebi.ac.uk/arrayexpress/). The TST transcriptome of the discovery cohort of patients with active TB was derived from dataset E-MTAB-3254, the cured TB TST transcriptome was derived from dataset E-MTAB-6815, and the validation active TB and latent TB TST transcriptomes were derived from dataset E-MTAB-6816.

### Immunostaining

Immunostaining of IL-17F was performed on 10 μm sections as previously described (51). Immunostaining of MMP-1 was performed as previously described (52). More detailed methods for both these targets, as well as approaches to quantify staining intensity, is available in Supplementary Methods.

## Acknowledgements

We thank our team of research nurses, in particular Cristina F Turienzo and Victoria Dean, for their assistance in recruiting patients and collecting samples, and the UCL-UCLH Pathogen Genomics Unit for RNASeq sample processing (https://www.ucl.ac.uk/infection-immunity/pathogen-genomics-unit). None of the authors have commercial affiliations or consultancies, stock or equity interests, or patent-licensing arrangements that could be considered a conflict of interest regarding the submitted manuscript.

