## Supplementary figures for "Exaggerated in vivo IL-17 responses discriminate recall responses in active TB"

**Supplementary Figure 1 (fig E1)**

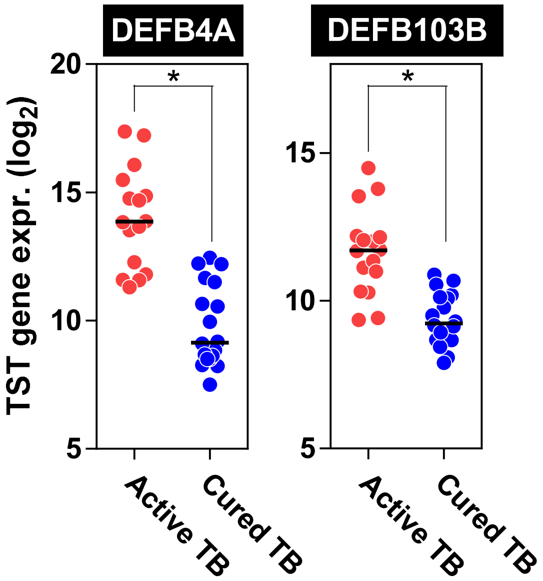


**Beta-defensin genes are enriched in active TB.**

mRNA expression of DEFB4A and DEFB103B genes in the TST of patients with active and cured TB. * = p<0.01 by Mann-Whitney test

**Supplementary Figure 2 (fig E2)**

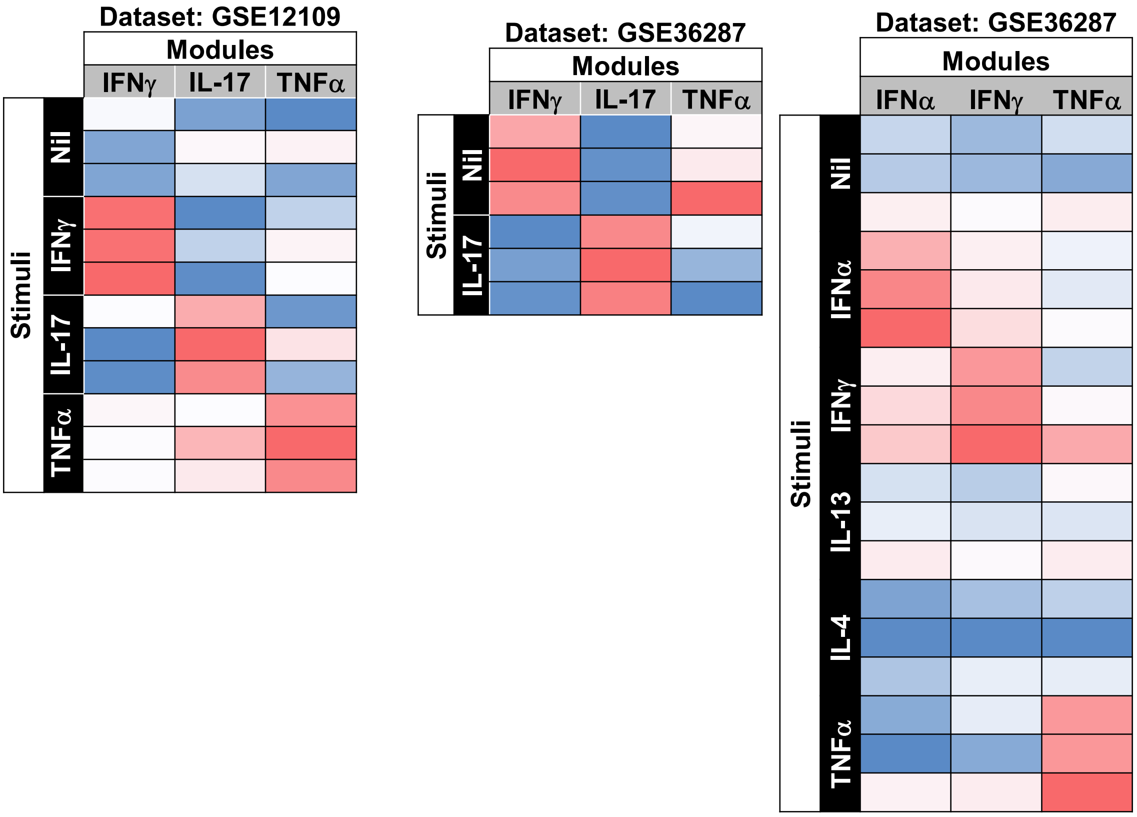


**Assessment of sensitivity and specificity of cytokine-stimulated keratinocyte transcriptional response modules.**

The expression of transcriptional modules derived from in vitro cytokine stimulation of keratinocytes (KC) determined in multiple datasets of in vitro stimulated KC.

Supplementary Figure 3 (fig E3)

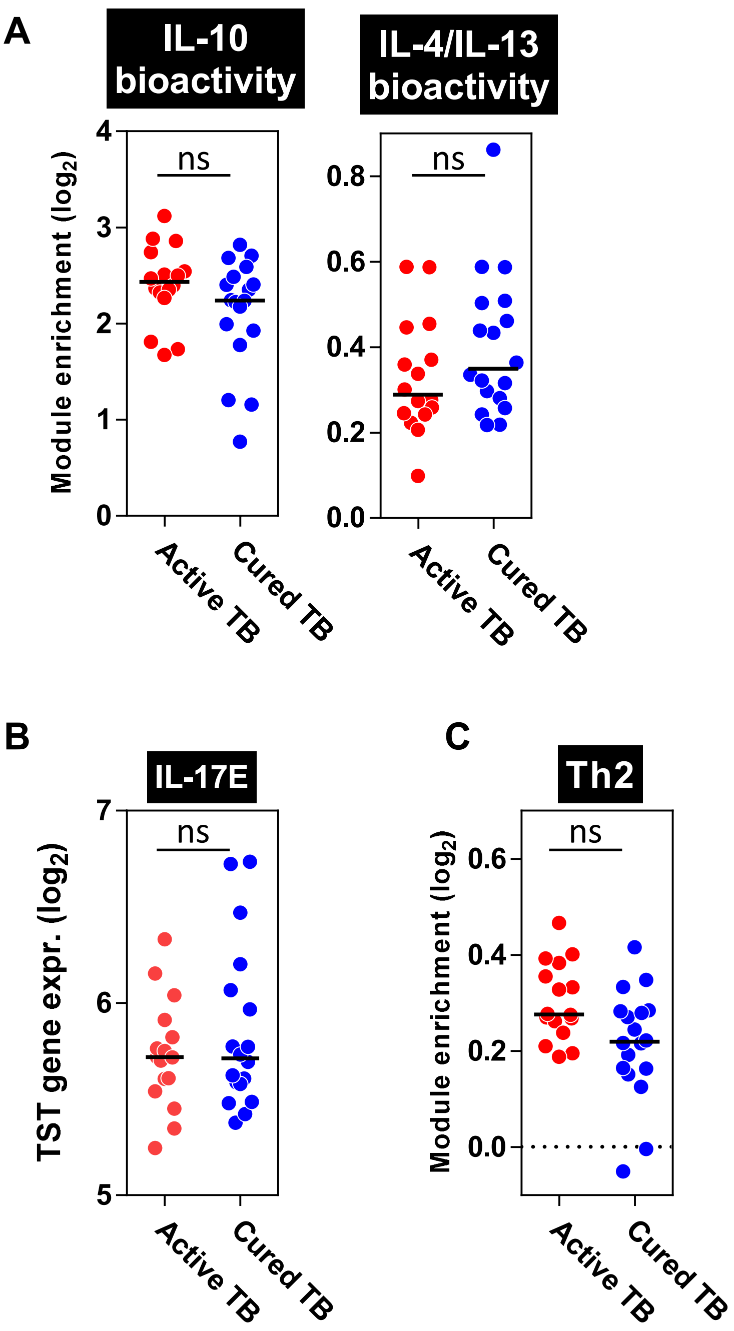


**Active TB is not characterised by IL-10 or Th2 immune responses.**

A) Enrichment in TST relative to saline injection of macrophage-response transcriptional modules. IL-10 and IL-4/IL-13 bioactivity modules derived from direct cytokine stimulation of macrophages. Module generation and validation available at Bell et al 2016 PloS Path. B) mRNA expression in TST of patients of IL17E gene. C) Enrichment in TST relative to saline injection of Th2 cell transcriptional module. All plots show no significant (ns) difference in module or gene expression between active and cured TB by Mann-Whitney test ($p>0.01$).

**Supplementary Figure 4 (fig E4)**

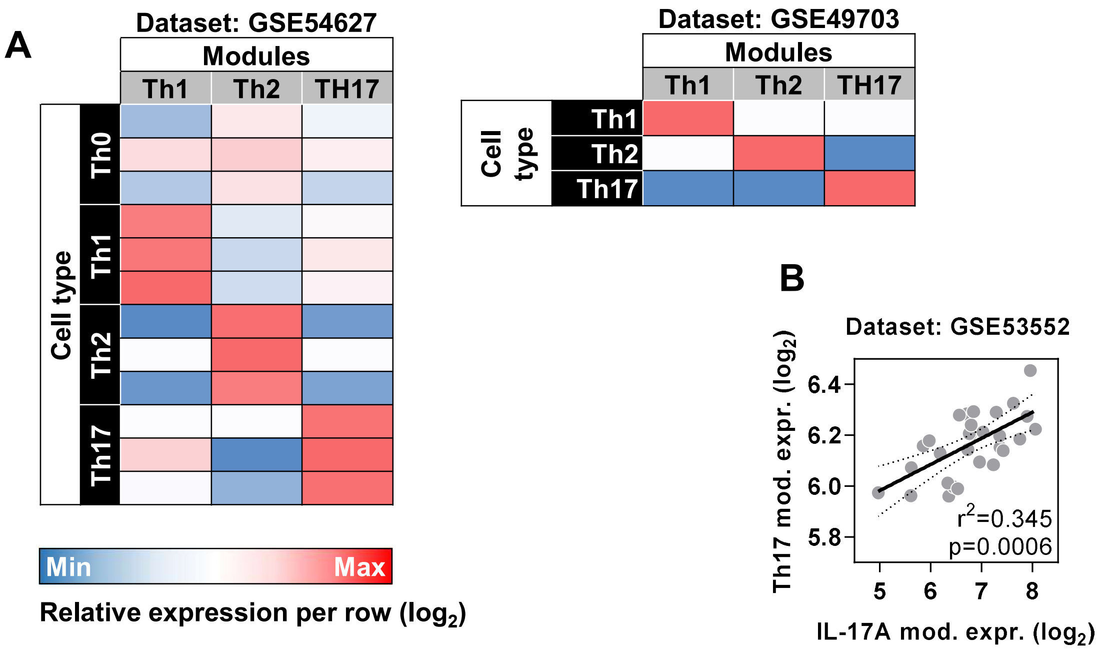


**Assessment of sensitivity and specificity of polarised CD4+ T helper cell transcriptional modules.**

**(A)** The expression of transcriptional modules derived from in vitro polarisation of CD4+ T cells in datasets containing T helper polarised phenotypes. **(B)** Relationship in the expression between IL-17A KC response module and Th17 module in the skin of patients with psoriasis vulgaris. r2 and p values determined by Spearman Rank Correlation.

**Supplementary Figure 5 (fig E5)**

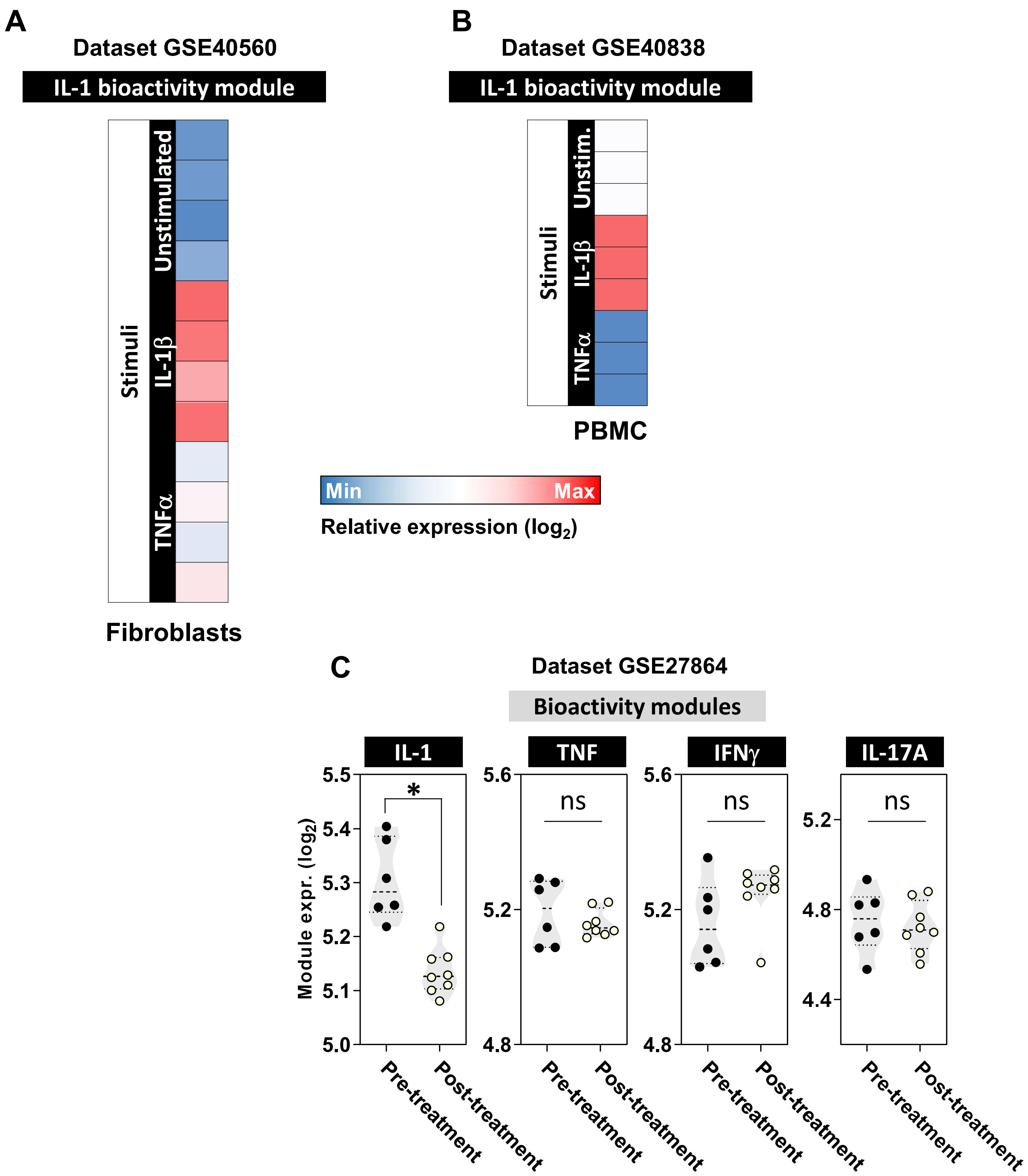


**Assessment of sensitivity and specificity of IL-1 transcriptional response module.**
The expression of IL-1 transcriptional response modules in A) fibroblasts stimulated in vitro for 6 hours with IL-1β (10 ng/ml) or TNFα (20 ng/ml), B) PBMC form healthy volunteers stimulated in vitro for 6 hours with IL-1β (10 ng/ml) or TNFα (20 ng/ml), and C) skin biopsies from Neonatal-onset Multisystem Inflammatory Disease patients before and after treatment with the IL-1 receptor antagonist, anakinra. * = $p<0.05$ by Mann-Whitney test. ns = non-significant.

**Supplementary Figure 6 (fig E6)**

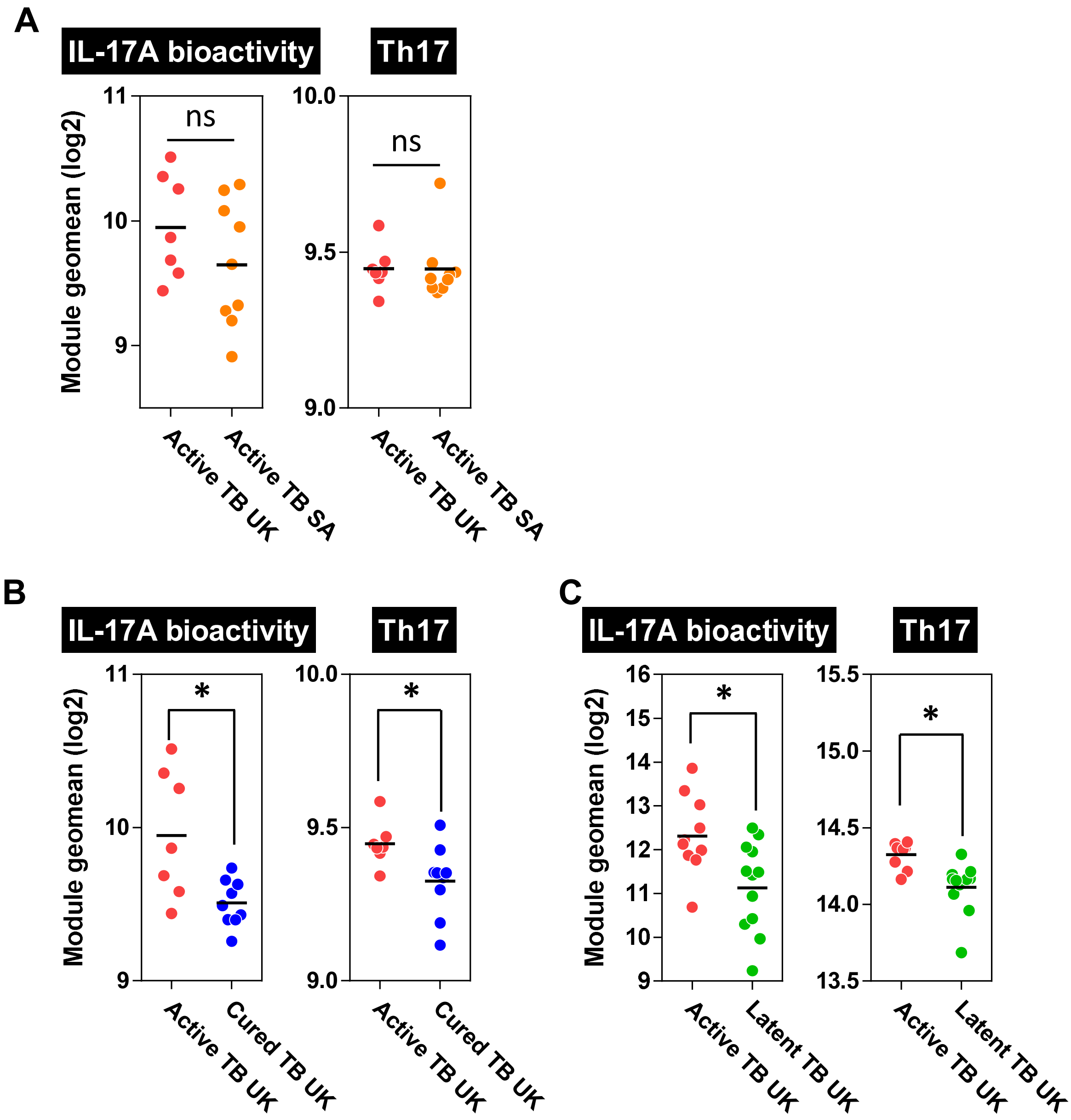


**IL-17 activity in TST in active TB is not confounded by demographic background.**

Geometric mean expression of IL-17A keratinocyte transcriptional response module and Th17 module in TST of A) active TB individuals with active TB recruited in the UK and South Africa, B+C) individuals with active and cured / latent TB recruited solely from the UK. * = p<0.05 by Mann-Whitney test. ns = non-significant.

**Supplementary Figure 7 (fig E7)**

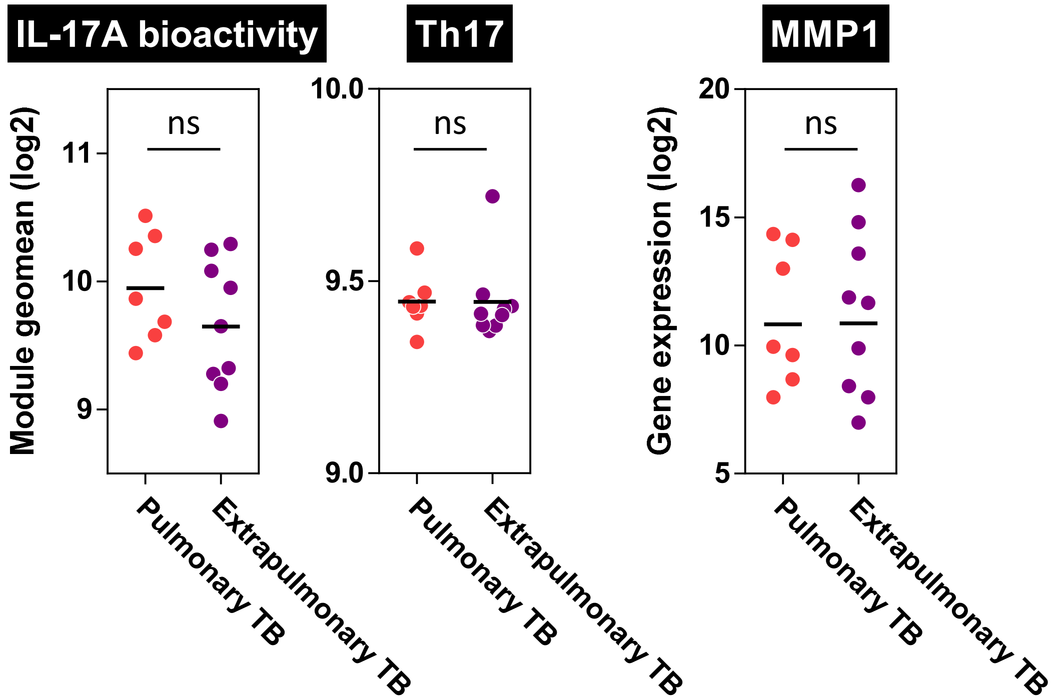


**IL-17A bioactivity, Th17 cell frequency and MMP-1 expression in TST in active TB is not confounded by extrapulmonary TB disease.**

Expression of IL-17A keratinocyte transcriptional response module, Th17 cell module and MMP1 gene mRNA in TST of active TB patients with pulmonary and extrapulmonary TB. ns = non-significant.

**Supplementary Figure 8 (fig E8)**

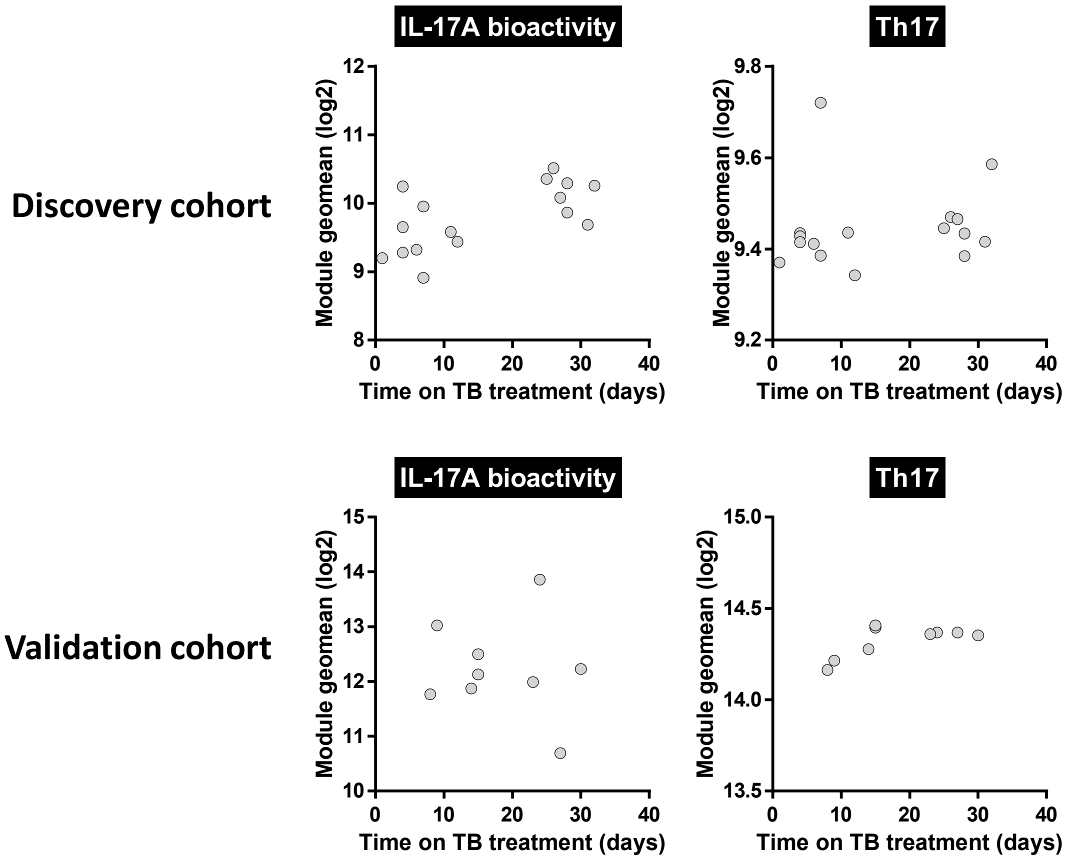


**IL-17A bioactivity and Th17 enrichment in TST in active TB is not confounded by time on treatment**

Relationship between duration of TB treatment at the time of TST sampling and geometric mean expression of either IL-17A keratinocyte transcriptional response or Th17 modules in TST. Assessment made in both discovery and validation cohorts of individuals with active TB disease.
