## Supplementary methods for "Exaggerated in vivo IL-17 responses discriminate recall responses in active TB"

#### Study populations

The study comprised recruitment of several different populations. The discovery ‘Active TB' group that formed the basis of most of the analyses in figs 1-5 was the HIV seronegative patients from the same cohort described in our previous publication (1), who were recruited from TB clinics in London, UK and Cape Town, South Africa, and were all within one month of commencing antibiotic therapy. The comparator ‘Cured TB’ group was an independent cohort recruited from TB clinics in London, UK and Lima, Peru who fulfilled the inclusion/exclusion criteria (table E1). All ‘Cured TB’ patients were less than 2 years after completion of curative anti-TB antibiotic therapy for drug sensitive disease. In addition, a separate validation cohort of patients with active TB was recruited from TB clinics in London (fig 6). These patients were also within one month of commencing antibiotic therapy. This population was compared to individuals with latent TB recruited from TB clinics in London and Lima, who fulfilled inclusion/exclusion criteria (table E1). All study participants were HIV seronegative and for those with active or cured TB, the presence of Mtb infection was confirmed by routine culture or molecular based methods according to local clinic protocols. The demographic, clinical and laboratory data for each study group is summarised in table E2.

#### Study approval

Recruitment of patients with cured and latent TB was approved by UK National Research Ethics Committee (reference number: 14/LO/0505) and Universidad Peruana Cayetano Heredia Institutional Ethics Committee (reference number: 62349). Recruitment of patients for the validation active TB cohort was approved UK National Research Ethics Committee (reference number: 16/LO/0776).

#### Study schedule and sampling

On recruitment to the study, all participants received 0.1 mL intradermal injection of two units tuberculin (Serum Statens Institute) or saline in the volar aspect of one forearm, and this site was marked with indelible ink. At 48 hours, the clinical response at the injection site was evaluated by measurement of the maximum diameter of inflammatory induration and two 3 mm adjacent punch biopsies were obtained from marked TST or saline injection site as previously described (2). One biopsy was placed in RNAlater (Thermo Fisher) and stored at -70°C, and the other biopsy was placed in 10%

formalin neutral buffered solution (Sigma-Aldrich) and stored at room temperature for at least 1 week prior to paraffin embedding.

#### Sample processing

The TST transcriptome from active TB patients in the discovery cohort was derived directly from the data repository E-MTAB-3254 (ArrayExpress - https://www.ebi.ac.uk/arrayexpress/). Skin samples from all other participants was stored in RNAlater at -70oC after collection. For processing, TST samples were equilibrated to room temperature for 30 minutes before being transferred to CK14 lysing kit tubes (Bertin Instruments) containing 350μl of Buffer RLT (Qiagen) supplemented with 1% 2-Mercaptoethanol (Sigma). Tubes were pulsed for 6 cycles on a Precellys Evolution homogeniser (Bertin Instruments), each cycle consisting of 23 seconds of homogenisation at speeds of 6300 rpm. Samples were rested on ice for 2 minutes between cycles. After homogenisation, cellular debris and lysing beads were precipitated by centrifugation and RNA isolated from the supernatant using RNeasy Mini Kit (Qiagen). Total RNA from TST samples of patients with cured TB was purified, labelled and hybridised on Agilent 8x60k microarrays as previously described (1). For the validation cohort of active TB and individuals with latent TB, the KAPA mRNA HyperPrep Kit (Roche Diagnostics) was used to construct stranded mRNA-Seq libraries from up to 500 ng intact total RNA after which paired-end sequencing was carried out using the 75 cycle high-output kit on the NextSeq 500 desktop sequencer (Illumina). Each run contained 24 samples and was demultiplexed using bcl2fastq by Illumina (https://support.illumina.com/sequencing/sequencing_software/bcl2fastq-conversion-software.html). Paired end reads were mapped to the Ensembl human transcriptome reference sequence (homo sapiens GRCh38, latest version available). Mapping and generation of read counts per transcript were done using Kallisto (3). R/Bioconductor package tximport was used to import the mapped counts data and summarise the transcripts-level data into gene level (4).

#### Whole genome transcriptional profiling and analysis software

Raw microarray data was processed and normalised as previously described (5). Raw RNASeq counts were normalised within-sample into TPM (transcripts per million) to remove feature-length and library-size effects (6). Log2 transformed TPM were used for further analysis. Data matrices from non-TST datasets were obtained from processed data series downloaded at the ArrayExpress repository. Probe identifiers were converted to gene symbols using platform annotations provided with each dataset. In circumstances where downloaded datasets were not log2 transformed, this was performed on the entire processed data matrix. Significant gene expression differences between datasets were calculated from normalised expression matrices using MultiExperiment Viewer v4.9 (http://www.tm4.org/mev.html). Pathway analysis was performed in InnateDB (7) and visualized as network diagrams in Gephi v0.8.2 beta. Upstream regulator analysis was performed using Ingenuity Pathway Analysis (Qiagen), focusing on cytokines with predicted activation z-score >2. Genes predicted to be regulated in either active or cured TB formed the basis of the network diagram in fig 3. The expression of transcriptional modules was determined by calculating the geometric mean expression of all the module constituent genes found in the dataset being analysed, using R scripts generated in our previous publication (8), and which are available to download and use from the Github repository (https://github.com/MJMurray1/MDIScoring). Venn diagrams were constructed using the BioVenn tool (http://www.biovenn.nl/).

#### Transcriptomic data repositories

All transcriptional datasets used in this study are described in table E5. Accession numbers refer to datasets in the ArrayExpress repository (https://www.ebi.ac.uk/arrayexpress/). The TST transcriptome of the discovery cohort of patients with active TB was derived from dataset E-MTAB-3254, the cured TB TST transcriptome was derived from dataset E-MTAB-6815, and the validation active TB and latent TB TST transcriptomes were derived from dataset E-MTAB-6816.

#### Module derivation and expression

Immune cell modules used were ones rated with the highest Module Discriminatory Index (MDI) score for module sensitivity and specificity as determined in our previous publication (8). These were “M19” (T cells), “M37.1” (neutrophils) and “Monocyte 2-fold” (monocytes). We also made use of another neutrophil module “ImSig” that was derived and validated elsewhere (9). We have previously generated and validated the specificity of macrophage response modules to IL-10 or IL4/IL-13 stimulation (1), and these are also utilised in this manuscript.

To derive keratinocyte (KC) cytokine-response modules, we made use of previously published transcriptomic data (GSE12109 & GSE36287) from primary human keratinocytes (KCs) stimulated with a selection of cytokines (10, 11). Significant transcriptional responses (paired t-test with $\alpha$ of $p<0.05$ and no multiple testing correction) of genes over-expressed >4-fold in the cognate cytokine condition relative to unstimulated KC were initially identified. Modules were then derived from genes that were not also upregulated > 2-fold by non-cognate cytokine conditions compared to unstimulated KC. The KC IL-17 response module utilised a cut-off of 2-fold between IL-17 stimulated KC and unstimulated

KC as too few genes were upregulated >4-fold compared to unstimulated KC. The constituent genes of the KC modules are shown in table E6. The KC TNFa response module generated in this way has already been published (12), but the other cytokine modules are newly described. The gene components of the KC cytokine-response modules are available in table E6, and their specificity was evaluated in both datasets in which they were derived and in independent datasets of in vitro cytokine-stimulated KC (fig E1A).

The transcriptome of CD4+ T cells polarised towards different T helper phenotypes was derived from dataset GSE54627 (13). To derive specific modules for each differentiation state, we used gene expression from cells stimulated with anti-CD3 and anti-CD28, identifying genes over-expressed in the phenotype of interest compared to all other conditions by paired t-test with $\alpha$ of $p<0.05$ and no multiple testing correction. Each module was derived from the unique genes over expressed by more than 1.5-fold in the cognate condition compared to all other stimulation conditions (except for Th1 module where a 2 -fold cut-off was used). The gene components of T helper modules are available in table E7, and their specificity was evaluated in the dataset from which they were derived, in an independent dataset of polarised CD4 T cells, and in skin biopsies of patients with psoriasis vulgaris (fig E1B) (13–15).

An IL-1β-response module was derived from dataset cytokine stimulated fibroblasts (16) using an unpaired t-test with $\alpha$ of $p<0.05$ without multiple testing correction, and identifying genes induced >2-fold in IL-1β stimulated fibroblasts compared to TNF-stimulated fibroblasts. The gene components of this IL-1β bioactivity module are available in table E8. The specificity of the module was evaluated in the dataset from which it was derived, in an independent dataset of IL-1β stimulated PBMC (Accession no: GSE40838) and in skin from individuals treated with recombinant IL-1 receptor antagonist, anakinra (Accession no: GSE27864) (65).

#### Immunohistochemistry

Immunostaining of IL-17F was performed on 10 μm sections as previously described (17). Briefly, monoclonal mouse-anti-IL-17F, 1:50, code MA5-16229 [Thermo Fisher Scientific, Waltham, MA, USA] and DAPI (sc-24941, Santa Cruz Biotechnology, Texas, Dallas, USA) for detecting nuclei were used. Irrelevant primary antibodies from Sigma (Milan, Italy; irrelevant mouse, 1:50, code I8765) were applied at the same concentration of the related specific primary antibodies for immunostaining of negative control slides, and as a further negative control, secondary antibody (AF568, goat anti-mouse; Invitrogen, Thermo Fisher Scientific, Waltham, MA, USA) alone was used (data not shown).

Light-microscopic analysis was performed at a magnification of 40x with a Leica DM4000B microscope equipped with DFC-320 Leica digital camera (Leica Microsystems, Wetzlar, Germany).

Immunostaining of MMP-1 was performed as previously described (18, 19). Scanned slide images were obtained with use of NanoZoomer Digital Pathology System (Hamamatsu, Japan. Quantification of MMP-1 staining was performed blindly by extracting MMP-1 associated 3, 3 -diaminobenzidine (DAB) stain using standard deconvolution protocols in ImageJ. Cellular infiltrates were manually selected as depicted in Fig 2 and DAB stain quantified by staining intensity as proportion of the area selected using ImageJ. The selected region was moved without resizing to adjacent tissue that did not contain cellular infiltration to calculate background MMP-1 intensity. Three cellular infiltrates and background tissue regions were analysed for each tissue samples. Six TST samples were quantified in each group.

### Supplementary methods references

10. Nograles KE, Zaba LC, Guttman E, Fuentes-Duculan J, Suarez-Farinas M, Cardinale I, Khatcherian A, Gonzalez J, Pierson KC, White TR, Pensabene C, Coats I, Novitskaya I, Lowes MA, Krueger JG. Th17 cytokines interleukin (IL)-17 and IL-22 modulate distinct inflammatory and keratinocyte-response pathways. *Br J Dermatol* 2008;159:1092–1102.
11. Swindell WR, Xing X, Stuart PE, Chen CS, Aphale A, Nair RP, Voorhees JJ, Elder JT, Johnston A, Gudjonsson JE. Heterogeneity of Inflammatory and Cytokine Networks in Chronic Plaque Psoriasis. *PLOS ONE* 2012;7:e34594.
12. Byng-Maddick R, Turner CT, Pollara G, Ellis M, Guppy NJ, Bell LCK, Ehrenstein MR, Noursadeghi M. Tumor Necrosis Factor (TNF) Bioactivity at the Site of an Acute Cell-Mediated Immune Response Is Preserved in Rheumatoid Arthritis Patients Responding to Anti-TNF Therapy. *Front Immunol* 2017;8:932.
13. Touzot M, Grandclaudon M, Cappuccio A, Satoh T, Martinez-Cingolani C, Servant N, Manel N, Soumelis V. Combinatorial flexibility of cytokine function during human T helper cell differentiation. *Nat Commun* 2014;5:.
14. Ramesh R, Kozhaya L, McKevitt K, Djuretic IM, Carlson TJ, Quintero MA, McCauley JL, Abreu MT, Unutmaz D, Sundrud MS. Pro-inflammatory human Th17 cells selectively express P-glycoprotein and are refractory to glucocorticoids. *J Exp Med* 2014;211:89–104.
15. Russell CB, Rand H, Bigler J, Kerkof K, Timour M, Bautista E, Krueger JG, Salinger DH, Welcher AA, Martin DA. Gene expression profiles normalized in psoriatic skin by treatment with brodalumab, a human anti-IL-17 receptor monoclonal antibody. *J Immunol* 2014;192:3828–3836.
16. Aubert P, Suárez-Fariñas M, Mitsui H, Johnson-Huang LM, Harden JL, Pierson KC, Dolan JG, Novitskaya I, Coats I, Estes J, Cowen EW, Plass N, Lee C-CR, Sun H-W, Lowes MA, Goldbach-Mansky R. Homeostatic Tissue Responses in Skin Biopsies from NOMID Patients with Constitutive Overproduction of IL-1β. *PLoS ONE* 2012;7:.

17. Ricciardolo FLM, Sorbello V, Folino A, Gallo F, Massaglia GM, Favatà G, Conticello S, Vallese D, Gani F, Malerba M, Folkerts G, Rolla G, Profita M, Mauad T, Di Stefano A, Ciprandi G. Identification of IL-17F/frequent exacerbator endotype in asthma. *J Allergy Clin Immunol* 2017;140:395–406.

18. Marafioti T, Paterson JC, Ballabio E, Reichard KK, Tedoldi S, Hollowood K, Dictor M, Hansmann M-L, Pileri SA, Dyer MJ, Sozzani S, Dikic I, Shaw AS, Petrella T, Stein H, Isaacson PG, Facchetti F, Mason DY. Novel markers of normal and neoplastic human plasmacytoid dendritic cells. *Blood* 2008;111:3778–3792.

19. Akarca AU, Shende VH, Ramsay AD, Diss T, Pane-Foix M, Rizvi H, Calaminici MR, Grogan TM, Linch D, Marafioti T. BRAF V600E mutation-specific antibody, a sensitive diagnostic marker revealing minimal residual disease in hairy cell leukaemia. *Br J Haematol* 2013;162:848–851.
